# Combination of cycling hyperthermia and Echinacoside creates synergistic curing effect on pancreatic cancer PANC-1 cells

**DOI:** 10.1101/2024.09.04.611320

**Authors:** Wei-Ting Chen, You-Ming Chen, Guan-Bo Lin, Yu-Yi Kuo, Hsu-Hsiang Liu, Chih-Yu Chao

## Abstract

Therapy targeting the suppression of human MutT homolog 1 (MTH1) has been gaining ground in recent years, thanks to its resulting significant increase of 8-hydroxy-2’-deoxyguanosine triphosphate (8-oxo-dGTP) accumulation in genomic DNA, causing DNA damage and apoptotic cell death. Echinacoside (Ech), a natural phenylethanoid glycoside first extracted from Echinacea angustifolia or desert plant Cistanches is one of a few natural products which are capable of inhibiting the MTH1 function. It, however, is difficult to apply it in clinical trials, due to high cost for effective dosage in need. In the study, we show that combination with thermal-cycling hyperthermia (TC-HT), a novel physical treatment, can amplify the curative effect of Ech, reducing its dosage in need significantly. The combination resulted in a multipronged mechanism targeting multiple key apoptotic regulating proteins such as Bcl-2 and MAPK family proteins. Its effect is expected to be comparable to the treatment strategy containing MTH1, Bcl-2, and ERK inhibitors, posing as new promising approach in cancer treatment.

## Introduction

Due to difficulty in early diagnosis, pancreatic cancer has remained a highly fatal disease with a five-year survival rate less than 10% in the U.S., and is predicted to be the second leading cause for cancer-related death in the next decade in Western countries. Despite progress in diagnosis and therapy, prognosis for pancreatic cancer patients has remained poor. Given the imperative need for progress in early diagnosis and therapeutic effect, it is worthwhile to seek novel anticancer agents and alternative therapies, in order to improve the life quality of pancreatic cancer patients.

Ingredients extracted from plants or traditional Chinese herbs, known as phytochemicals, have emerged as promising novel anticancer drug sources, on top of their application in anti-virus, anti-cancer and anti-inflammatory therapies (1–4). A main reason for growing popularity of herbal drugs is their absence of side effects. Experiments have shown that anticancer phytochemicals attain curative effect via regulating molecular pathways implicated in proliferation and progression of cancer cells. A noticeable example is Echinacoside (Ech), a natural phenylethanoid glycoside extracted from Echinacea angustifolia or desert plant Cistanches. Ech has been proven to be capable of rescuing the PC-12 neuronal cells from neurotoxicity via attenuating mitochondrial dysfunction and inflammatory responses by reducing ROS production (5). Recent studies have also shown that Ech can improve the behavioral disorders in mouse model with Parkinson’s disease (6). Besides, the hepatoprotection and anti-senescence effects of Ech have also been found in mouse models (7). However, the specific mechanism responsible for Ech’s anticancer effect remains unknown, as only a few cancer cell lines have been studied (8–10).

Recent studies have manifested that Ech is capable of inhibiting the catalytic activities of MutT homolog 1 (MTH1), a protein responsible for sanitizing oxidized deoxynucleoside triphosphate (dNTP), highly expressive in various cancer cells (11). According to recent studies (12,13), MTH1 also plays a key role in the survival of cancer cells, helping them to avoid incorporation of oxidized dNTPs, which cause DNA damage and cell death. Noticeably, the inhibition of MTH1 doesn’t affect the growth and survival of normal cells (14), underscoring the potential of such therapeutic approach. Some MTH1 inhibitors have been developed with the aim of suppressing cancer growth by damaging DNA in cancer cells (15). In the study, we aimed to explore the anticancer effect of combining Ech with thermal treatment and the underlying mechanism.

As an adjuvant therapy, Hyperthermia (HT) has been applied along with radiation or chemotherapy attaining synergistic curative effect for cancer patients (16). However, heat generated by continuous HT therapy may damage normal cells and cause other side effects, due to lack of precision and flexible control of heating temperatures and parameters. To cope with the problem, our previous study proposed a novel thermal therapy, namely thermal cycling hyperthermia (TC-HT), for the anticancer treatment in pancreatic cancer cells (17,18). The TC-HT treatment, which features alternating application of high and low temperatures in combination with administration of various natural compounds, has significantly enhanced curative effect without harming normal cells.

In this paper, the study proposes an innovate treatment for pancreatic PANC-1 cancer cells, combining the natural compound Ech and the physical stimulus TC-HT, both of which are capable of inhibiting the growth of cancer cells with minimum injury to normal cells. Our results showed that TC-HT alone can inhibit the expression level of MTH-1 protein significantly. Moreover, the combined treatment with Ech demonstrated remarkable synergistic effect on inhibition of the expression level of MTH-1, with TC-HT amplifying Ech-induced apoptotic cell killing and thus reducing its necessary dosage. The remarkable anticancer performance could be attributed to TC-HT enhancing the pharmacological effect of Ech to inhibit MTH1 expression and cause 8-hydroxy-2’-deoxyguanosine triphosphate (8-oxo-dGTP) accumulation, thereby resulting in DNA damage and cell apoptosis. The combination treatment of Ech and TC-HT also significantly down-regulated ERK and Bcl-2 proteins, which are associated with cell proliferation and survival. The approach, thus, can attain an anticancer effect comparable to chemotherapy drugs with minimum damage to normal cells, shedding light on the potential of other combination therapies.

## Materials and methods

### Cell culture and treatment

The human pancreatic cancer PANC-1 cell lines were purchased from the Bioresource Collection and Research Center (BCRC, Hsinchu, Taiwan) and maintained in DMEM (HyClone; Cytiva, Marlborough, MA, USA) supplemented with 10% fetal bovine serum (HyClone; Cytiva) and 1% penicillin-streptomycin (Gibco; Thermo Fisher Scientific, Inc., Waltham, MA, USA). H6c7 human pancreatic duct epithelial cell line was obtained from Kerafast, Inc. (Kerafast, Inc.; Absolute Biotech, Boston, MA, USA) and maintained in keratinocyte-serum free medium (Invitrogen; Thermo Fisher Scientific, Inc., Carlsbad, CA, USA) supplemented with human recombinant epidermal growth factor, bovine pituitary extract (Invitrogen; Thermo Fisher Scientific, Inc.), and 1% (v/v) penicillin and streptomycin. All cells were maintained in a humidified incubator with 5% CO_2_ and 95% air at 37°C. Ech (Sigma-Aldrich; Merck KGaA, Darmstadt, Germany) was dissolved in distilled water at a concentration of 2 mg/ml. Cells were plated in 24-well plates or 3-cm culture dishes 24 h before treatment with or without TC-HT and/or Ech. In the experiment, after Ech was added to cells, TC-HT treatment was applied using the Thermal Cycler (Applied Biosystems; Thermo Fisher Scientific, Inc., Waltham, MA, USA). The actual temperatures of cancer cells were measured by a needle thermocouple located in the bottom of the well. After the treatment, cells were incubated in a cell culture incubator until further analysis.

### Cell viability assay

After the treatment, cell viability was determined by 3-(4,5-dimethylthiazol-2-yl)-2,5-diphenyltetrazolium bromide (MTT) (Sigma-Aldrich; Merck KGaA) assay. The medium was replaced with MTT solution (0.5 mg/ml in DMEM) and incubated at 37°C for 4 h. Then the supernatants were discarded, and the remaining formazan crystals were dissolved with dimethyl sulfoxide and the optical density in each well was then evaluated by the measurement of absorbance at 570 nm using an ELISA microplate reader. The cell viability was calculated based on the optical intensity of the formazan, and was expressed as a percentage of the untreated controls, which were set at 100%.

### Flow cytometric detection of apoptotic cells

Apoptotic cells were examined using an Annexin V-FITC and propidium iodide (PI) double-staining kit (BD Biosciences, Franklin Lakes, NJ, USA) by flow cytometry. Cells used for flow cytometry were collected by trypsinization, washed twice with PBS, and resuspended in binding buffer containing Annexin V-FITC and PI. Cells were stained for 15 min at room temperature in the dark before being analyzed by flow cytometry.

### ROS level detection

Intracellular ROS levels were measured by flow cytometry using the fluorescent dye dihydroethidium (DHE) (Sigma-Aldrich; Merck KGaA). PANC-1 cells were harvested 24 h after treatments and rinsed with PBS before staining. Cells used for flow cytometry were collected by trypsinization and then incubated with 5 µM DHE for 30 min at 37°C in the dark. The fluorescence intensity emitted from DHE was measured by flow cytometry, and the ROS levels were expressed as mean fluorescence intensity for comparison.

### Measurement of intracellular 8-oxo-dGTP

It is well known that Avidin binds with high specificity to 8-oxo-dGTP and used to evaluate the incorporation of 8-oxo-dGTP into DNA. To observe cells under a confocal microscope, they were seeded on coverslips. Cells treated with Ech and/or TC-HT on coverslips were fixed with ice-cold methanol for 20 min followed by incubation in Tris-buffered saline (TBS) with 0.1% Triton X-100 for 15 min. Cells were blocked in 15% FBS, 0.1% Triton X-100 in TBS for 2 h at room temperature. Then, the cells were probed with Alexa Fluor 488-conjugated Avidin (Alexa 488) (Invitrogen; Thermo Fisher Scientific, Inc.) in blocking solution for 1 h at 37 °C. After washing with TBS containing 0.1% Triton X-100 for three times, the coverslips were mounted on glass slides in mounting medium with DAPI (Abcam, Cambridge, United Kingdom). Fluorescent images were taken by a Zeiss LSM 880 inverted laser scanning confocal microscope. These images were then analyzed with ImageJ v3.91 software to calculate the relative fluorescence units (RFU) for both green and blue fluorescence. The level of 8-oxo-dGTP in cells was determined by normalizing the RFU ratio of green (Alexa 488) to blue (DAPI) fluorescence against the control group.

### Western blot analysis

The levels of protein expression in PANC-1 cells were examined by Western blot analysis. After treatment with Ech or TC-HT or in combination, cells were scraped from culture dishes and lysed in ice-cold RIPA lysis buffer containing protease inhibitor (EMD Millipore, Billerica, MA, USA). The lysates were centrifuged at 12,000 × g for 30 min at 4°C, and the supernatants were collected and the protein concentrations were determined by Bradford protein assay (BioShop Canada Inc., Burlington, Canada). Equal amount of proteins (30 µg) were resolved on 10% SDS-PAGE and then transferred onto polyvinylidene fluoride membranes (EMD Millipore). Nonspecific antibody binding sites were blocked in 5% nonfat milk in TBS with Tween-20 (TBST; 20 mM Tris-base, pH 7.6; 0.15 M NaCl; and 0.1% Tween-20) for 1 h at room temperature. The blocked membranes were probed with MTH1, p-ERK, t-ERK, p-JNK, Bcl-2, Bax, PARP (all from Cell Signaling Technology, Inc., Danvers, MA, USA) and GAPDH (GeneTex, Inc., Irvine, CA, USA) antibodies overnight at 4°C. The washed membranes were then incubated with horseradish peroxidase-conjugated goat anti-rabbit secondary antibodies (Jackson ImmunoResearch Laboratories, Inc., West Grove, PA, USA) in a blocking solution. Immunoreactivity signal was visualized with an enhanced chemiluminescence substrate (Advansta, Inc., Menlo Park, CA, USA) and detected by the Amersham Imager 600 imaging system (GE Healthcare Life Sciences, Chicago, IL, USA). Image analysis was performed with Image Lab software (Bio-Rad Laboratories, Inc., Hercules, California, USA).

### Statistical analysis

Statistical analyses were performed by using the OriginPro 2015 software (OriginLab). The results were expressed as the mean ± standard deviation, and each data point represented the average from three independent experiments. Differences of statistical significance were determined by one-way ANOVA followed by Tukey’s post-hoc test. P < 0.05 was considered to indicate a statistically significant difference.

## Results

### TC-HT for in vitro application

The detailed procedure of TC-HT treatment was described in our previous work (17). Briefly, TC-HT was adopted to perform a thermal cycling treatment with a high temperature period for 3 min and a cooling period for 30 sec, and this protocol was repeated continuously for ten cycles. In the study, the temperature variation of the cells under TC-HT was controlled by a modified PCR machine. The actual temperature in the culture well was measured by a needle thermocouple. Fig. 1 demonstrates the temperature variation sensed by the cells at the bottom of the culture well during TC-HT treatment.

**Figure 1.**
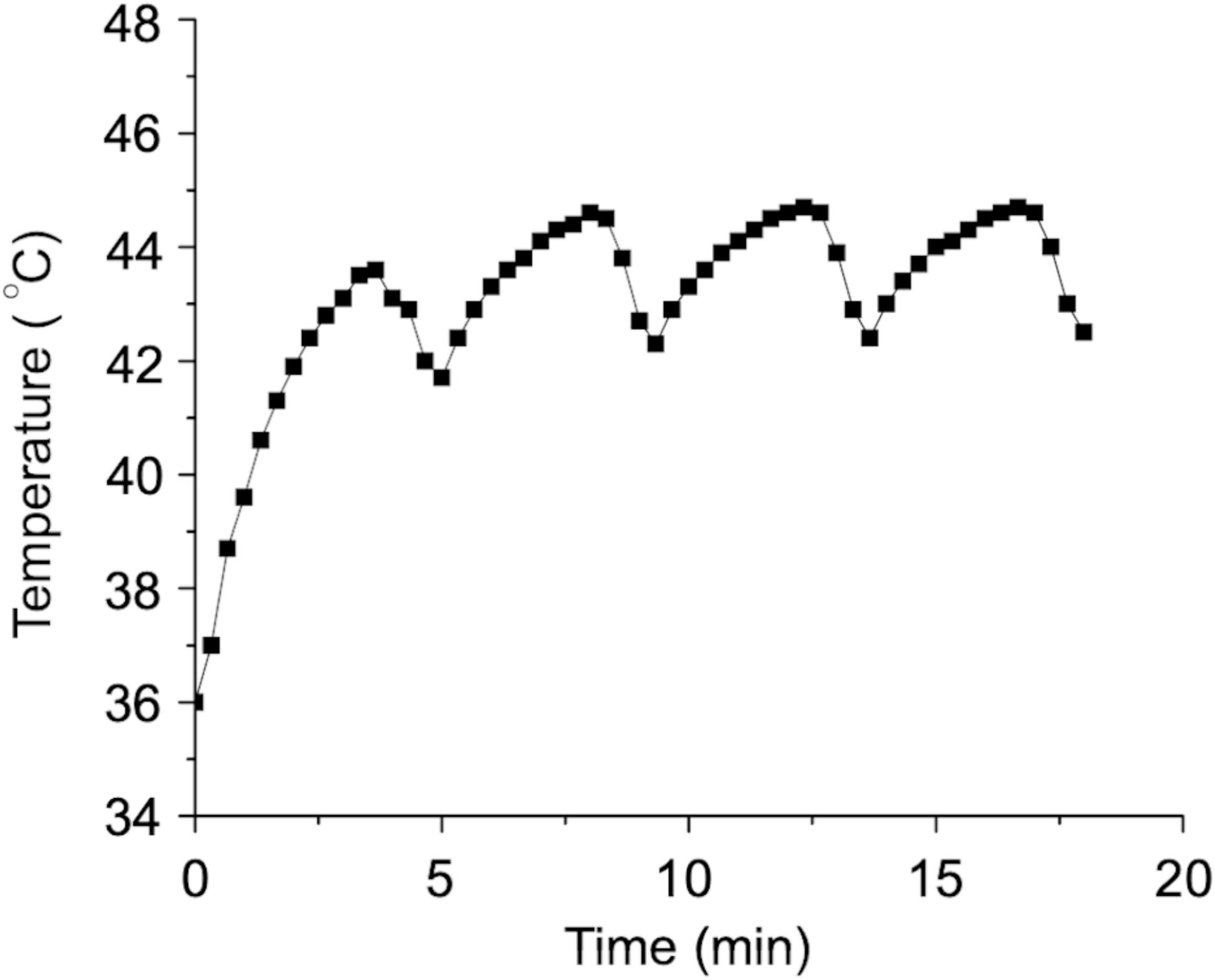
Temperature variation of TC-HT treatment. The temperature was raised to a desired high temperature for 3 min and decreased to a lower temperature for 30 s. The high and low temperature periods of TC-HT were repeated continuously for ten times in this study. The actual temperature in the culture well was monitored every 20 seconds by a needle thermocouple.

### Ech and TC-HT synergistically suppress PANC-1 cell proliferation

To examine the inhibitory effect of TC-HT and Ech, the viability of PANC-1 cells under TC-HT and/or Ech administration was evaluated by the MTT assay. As shown in Fig. 2A, Ech decreased the cell viability of PANC-1 cells in a concentration-dependent manner. It was found that 20 µM Ech treatment on PANC-1 cells reduced the cell viability to 80% compared to the control cells, which was used in the following experiments. In this study, we aim to investigate the combination anticancer effect of Ech and TC-HT, and thus low-dose Ech (20 µM) with mild inhibitory effect was adopted in the combination treatment. The results also showed that TC-HT alone only caused slight inhibitory effect as well as low-dose Ech. Moreover, we found that combining TC-HT and Ech greatly decreased the viability of PANC-1 cells to only 29% of the control group at the 20 µM low-dose Ech administration. Noteworthily, the inhibitory effect of low-dose Ech (20 µM) in combination with TC-HT was close to that of 100 µM Ech, indicating that the anticancer effect was highly amplified and therefore the inhibitory concentration of Ech combined with TC-HT could be decreased to one fifth of the Ech treatment alone. On the other hand, the H6c7 human pancreatic duct epithelial cell line was treated with 20 µM Ech alone or in combination with TC-HT to verify the toxic effect of the combination treatment on normal pancreatic cells. As shown in Fig. 2B, the combination treatment did not cause significant impact on the cell viability of the H6c7 normal cells, while it exerted significant apoptotic cell death on PANC-1 cancer cells in similar condition.

**Figure 2.**
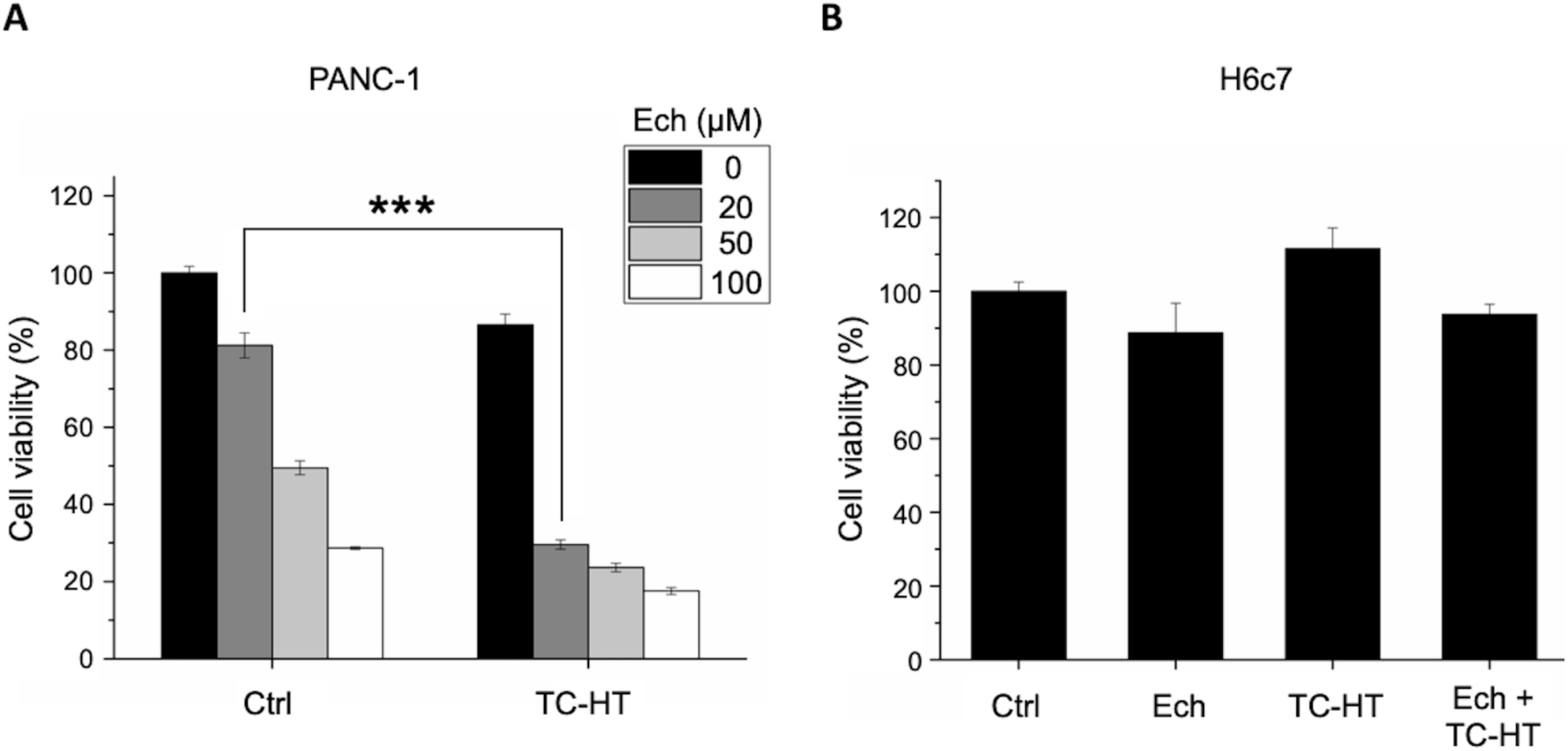
Effect of Ech alone or in combination with TC-HT on the cell viability. (A) MTT viability assay of PANC-1 cells treated with different concentrations of Ech or in combination with TC-HT treatment. (B) MTT viability assay of H6c7 normal human pancreatic cells treated with 20 µM Ech or in combination with the same TC-HT treatment. Data represent the mean ± standard deviation (n=3). Statistical significance was determined by one-way ANOVA followed by Tukey’s post-hoc test (***P < 0.001).

### Ech and TC-HT synergistically induce apoptosis in PANC-1 cells

To confirm whether the decreased cell viability was associated with the induction of apoptosis, flow cytometry with Annexin V-FITC and propidium iodide (PI) double staining was performed. Annexin V is a phospholipid binding protein which has a strong affinity for phosphatidylserine (PS) residues, and therefore can be used as a probe for detecting apoptosis. PI, a membrane-impermeable nucleic acid dye, was used to distinguish between the necrotic and apoptotic cells. The dot plot results (Fig. 3A) show a remarkable increase of apoptotic cell numbers (Q2+Q4) in the combination treatment group. The bar graph (Fig. 3B) shows that the average apoptotic rate of the combination treatment group is significantly higher than that treated with Ech or TC-HT alone. It demonstrated that co-treatment with TC-HT could synergize with Ech in inhibiting the growth of PANC-1 cancer cells through inducing cell apoptosis.

**Figure 3.**
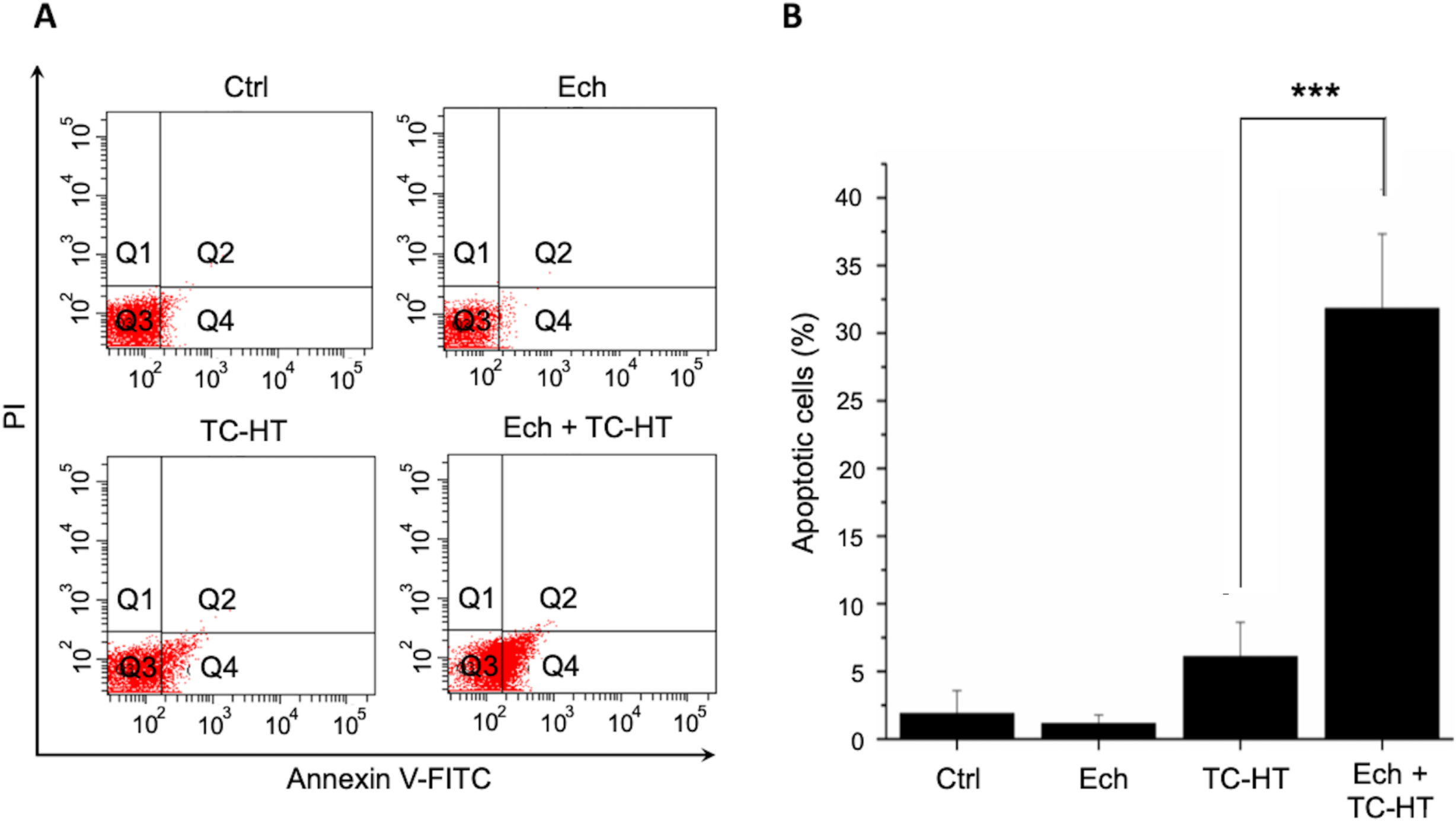
Assessment of the mode of cell death via Annexin V-FITC/PI double staining. (A) PANC-1 cells were treated with 20 µM Ech or in combination with TC-HT. After 24 h, the apoptotic state was assessed by flow cytometry with Annexin V/PI double staining. (B) The quantified bar graph showed that the percentage of apoptotic cells (Q2+Q4) was significantly higher in the combination treatment group. Data represent the mean ± standard deviation (n=3). Statistical significance was determined by one-way ANOVA followed by Tukey’s post-hoc test (***P < 0.001).

### Ech and TC-HT synergistically decrease MTH1 and increase cellular ROS levels

Previous study reported that Ech could serve as an MTH1 inhibitor by enzyme catalyzed reaction (11). In this study, we verify the effect of Ech treatment on MTH1 protein expression by Western blot analysis. As shown in Fig. 4A, the Western blot result demonstrated that 20 µM Ech treatment slightly reduced the MTH1 protein expression. Moreover, we found that TC-HT alone significantly decreased the expression level of MTH1 protein, while the combination of Ech and TC-TH further inhibited the expression of MTH1 protein synergistically. In cancer cells, MTH1 is known as a key factor for the remediation of ROS-mediated DNA damage caused by 8-oxo-dGTP.

**Figure 4.**
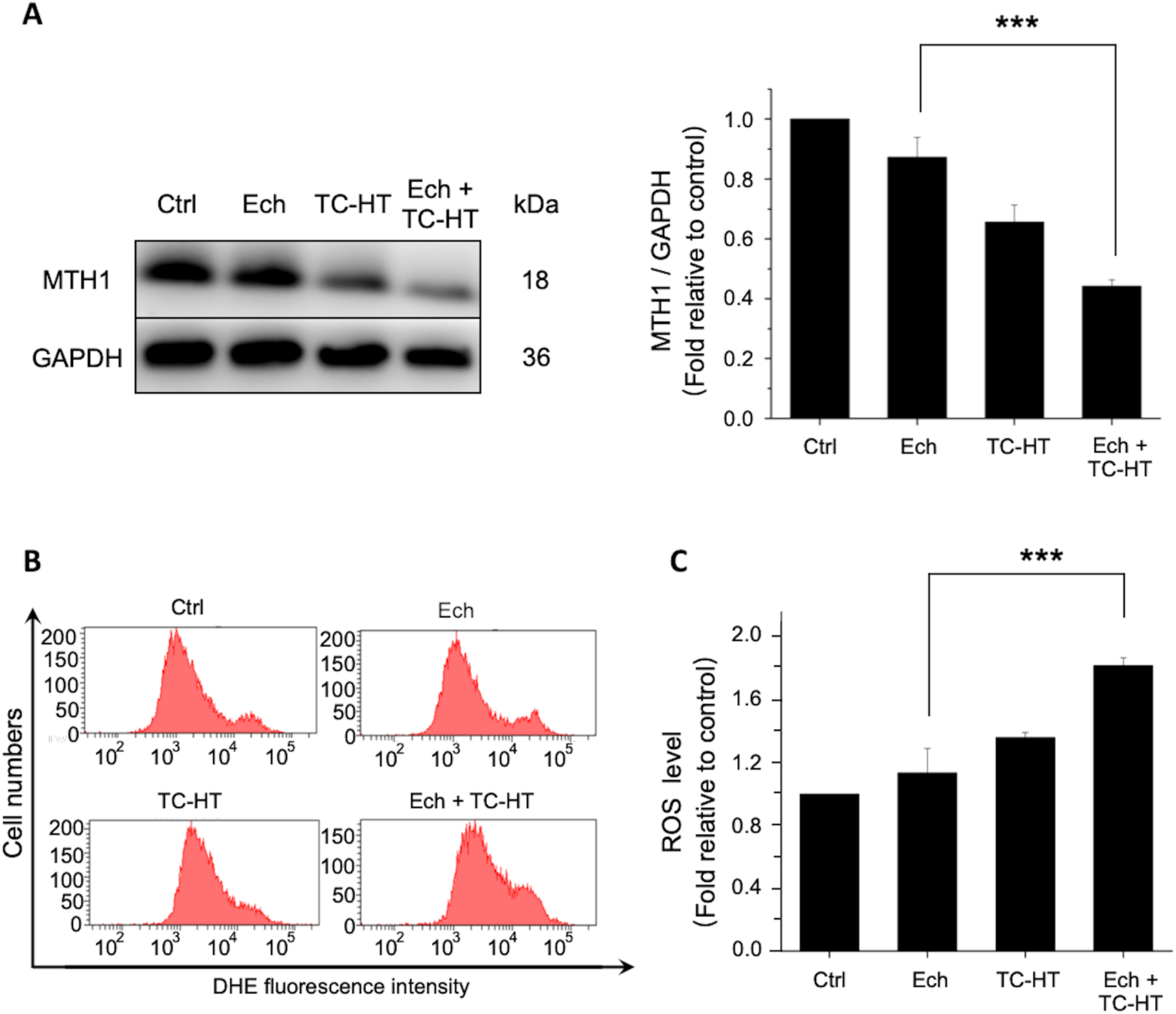
Combined effect of Ech and TC-HT on MTH1 protein expression and ROS level in PANC-1 cells. PANC-1 cells were treated with 20 µM Ech or in combination with TC-HT. After 24 h, the MTH1 protein expression was quantified by Western blot analysis and ROS level was assessed by flow cytometry. (A) The MTH1 protein expression was quantified and normalized to GAPDH internal control. (B) The ROS level was assessed by DHE fluorescent dye, and the flow cytometry result was shown as fluorescence intensity versus cell numbers. (C) The mean fluorescence intensity was quantified as bar graph representing the ROS level. Data represent the mean ± standard deviation (n=3). Statistical significance was determined by one-way ANOVA followed by Tukey’s post-hoc test (***P < 0.001).

To examine if cellular ROS level was affected by Ech and TC-HT co-treatment, the fluorescent dye DHE was used to detect the ROS level. As it can be seen from Fig. 4B and Fig. 4C, treatment with Ech alone did not significantly increase the ROS level in PANC-1 cells. On the other hand, the results showed that the treatment with TC-HT or the combination treatment with Ech and TC-HT significantly increased the ROS level in PANC-1 cells. In addition, it is noteworthy that the combination treatment (Ech + TC-HT) induced the most significant ROS level elevation, which may result in a large amount of DNA damage in cancer cells. The results suggest that Ech and TC-HT can work synergistically to increase the cellular ROS level in PANC-1 cells, causing the 8-oxo-dGTP accumulation to damage genomic DNA.

### Ech and TC-HT synergistically increase accumulation of intracellular 8-oxo-dGTP

Since 8-oxo-dGTP is the major oxidized nucleotide inside cells, which tends to be incorporated into DNA during replication, it leads to severe genome mutations and cell death consequently. Cellular 8-oxo-dGTP is generated by ROS, therefore, increase in cellular ROS would also result in increase in cellular 8-oxo-dGTP level. To directly verify that more DNA mutants caused by 8-oxo-dGTP were accumulated in the cells by the combination treatment (Ech + TC-HT), Alexa 488 was used to stain PANC-1 cells after the treatment. Since avidin has been shown to bind to 8-oxo-dGTP with high specificity, the DNA mispairing can thus be visualized by the green fluorescence in the immunofluorescence images (14). Fig. 5A shows the immunofluorescence increased in PANC-1 cells treated with 20 µM Ech or TC-HT for 24 hours, and significantly increased in cells treated with the combination of TC-HT and 20 µM Ech. The normalized RFU ratios of green/blue fluorescence (Fig. 5B) reveal that cells treated with the combination of Ech and TC-HT presented the highest 8-oxo-dGTP levels among the four groups.

**Figure 5.**
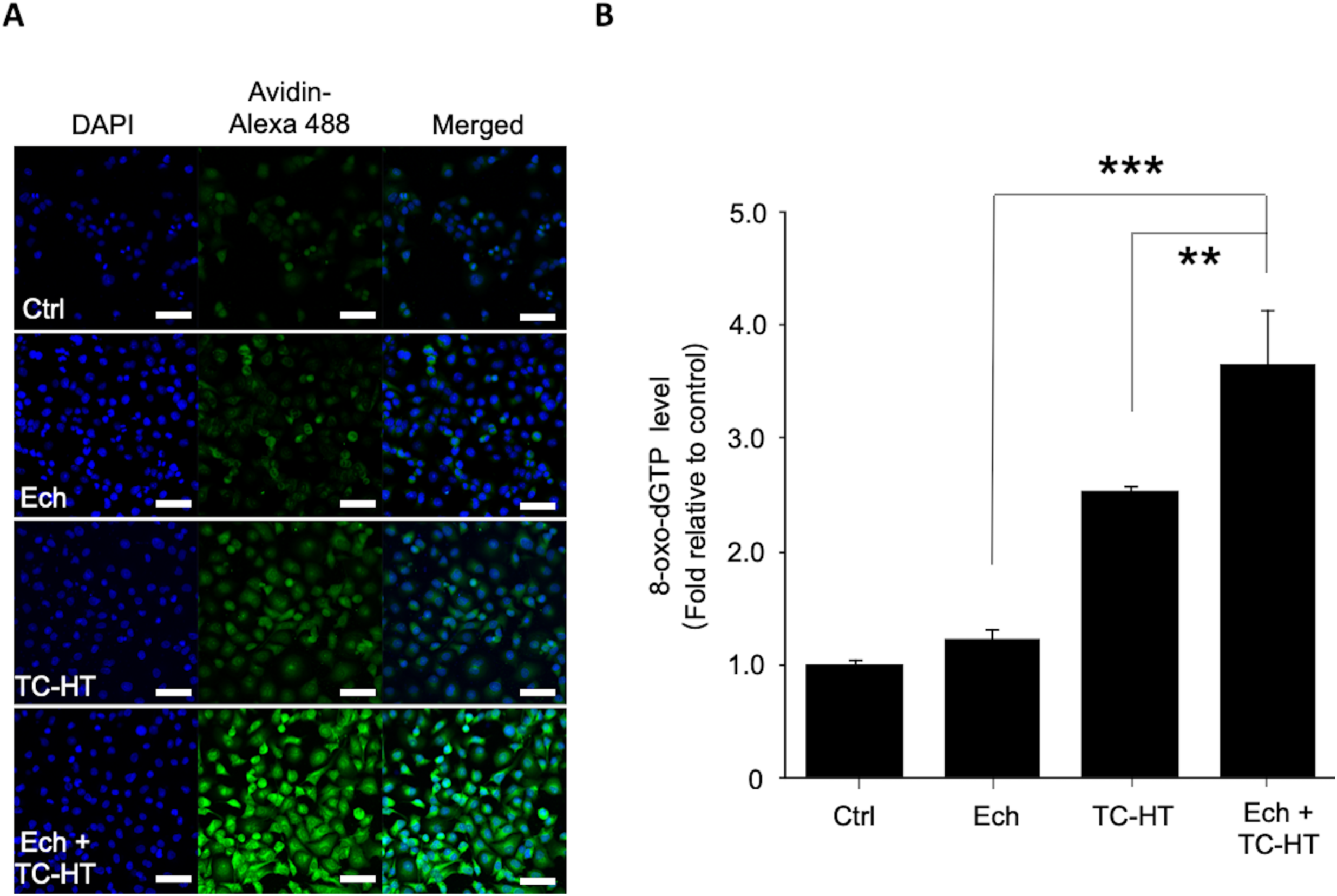
Enhanced accumulation of 8-oxo-dGTP in PANC-1 cells with Ech and TC-HT combination treatment. PANC-1 cells were treated with 20 µM Ech alone or in combination with TC-HT. After 24 h, the fluorescent images were taken by an inverted laser scanning confocal microscope. (A) The confocal immunofluorescence micrographs contained DAPI (blue), Alexa 488 (green), and the colocalization of both stainings. The merged image shows Alexa 488 staining for DNA damage in green and DAPI staining for nuclei in blue, indicating areas where DNA damage occurs within the nuclei. Scale bar = 50 µm. (B) The 8-oxo-dGTP levels of cells were quantified by calculating the normalized RFU ratio of green/blue fluorescence (Alexa 488/DAPI) relative to the control group. Data represent the mean ± standard deviation (n=3). Statistical significance was determined by one-way ANOVA followed by Tukey’s post-hoc test (***P < 0.001).

### TC-HT enhances the anticancer effect of Ech via modulating the MAPK family proteins and inducing the apoptosis pathway in PANC-1 cells

To further investigate the mechanism of TC-HT enhanced anticancer effect, the expression levels of the MAPK family proteins and apoptosis-related proteins were examined using Western blot analysis. There are three major subfamilies of MAPK proteins which have been clearly characterized to play important role in cellular functions: namely ERK, JNK, and p38 proteins (19). The MAPK cascades are responsible to relay, amplify and integrate extracellular signals to cellular responses such as proliferation, differentiation, transformation, and apoptosis. It is known that ERK proteins are important for cell survival and found to be upregulated in human tumors, which led to the development of ERK inhibitors for cancer therapeutics (20). JNK and p38, on the other hand, were deemed as stress responsive proteins and thus involved in cell apoptosis (21). Therefore, inhibition of ERK and concurrent activation of JNK and p38 are critical for inducing apoptosis (22). In this study, the results show that p-JNK (Fig. 6A) was strongly activated in the combination treatment of Ech and TC-HT, while p-ERK (Fig. 6B), on the other hand, was significantly inhibited. In addition, the Bcl-2 family proteins are key regulators of the mitochondria pathway of apoptosis, where the Bax/Bcl-2 ratio can be used to assess the upregulation of the apoptotic signal. As shown in Fig. 6C, we found that the Bax/Bcl-2 ratio was significantly elevated in the combined treatment group, suggesting that TC-HT and Ech could synergistically trigger the mitochondria-dependent apoptosis pathway. Moreover, poly (ADP-ribose) polymerase (PARP) serves as an important nuclear enzyme in DNA repairing. The cleavage of PARP by executioner caspases is the hallmark feature of apoptosis, which leads to DNA damage. In this study, the cleaved PARP level was examined by Western blot analysis. The results showed that co-treatment of Ech and TC-HT greatly increased the expression of cleaved PARP proteins compared to the control and single treatment groups (Fig. 6D), indicating that the combination of Ech and TC-HT synergistically activated caspase-dependent apoptosis pathway.

**Figure 6.**
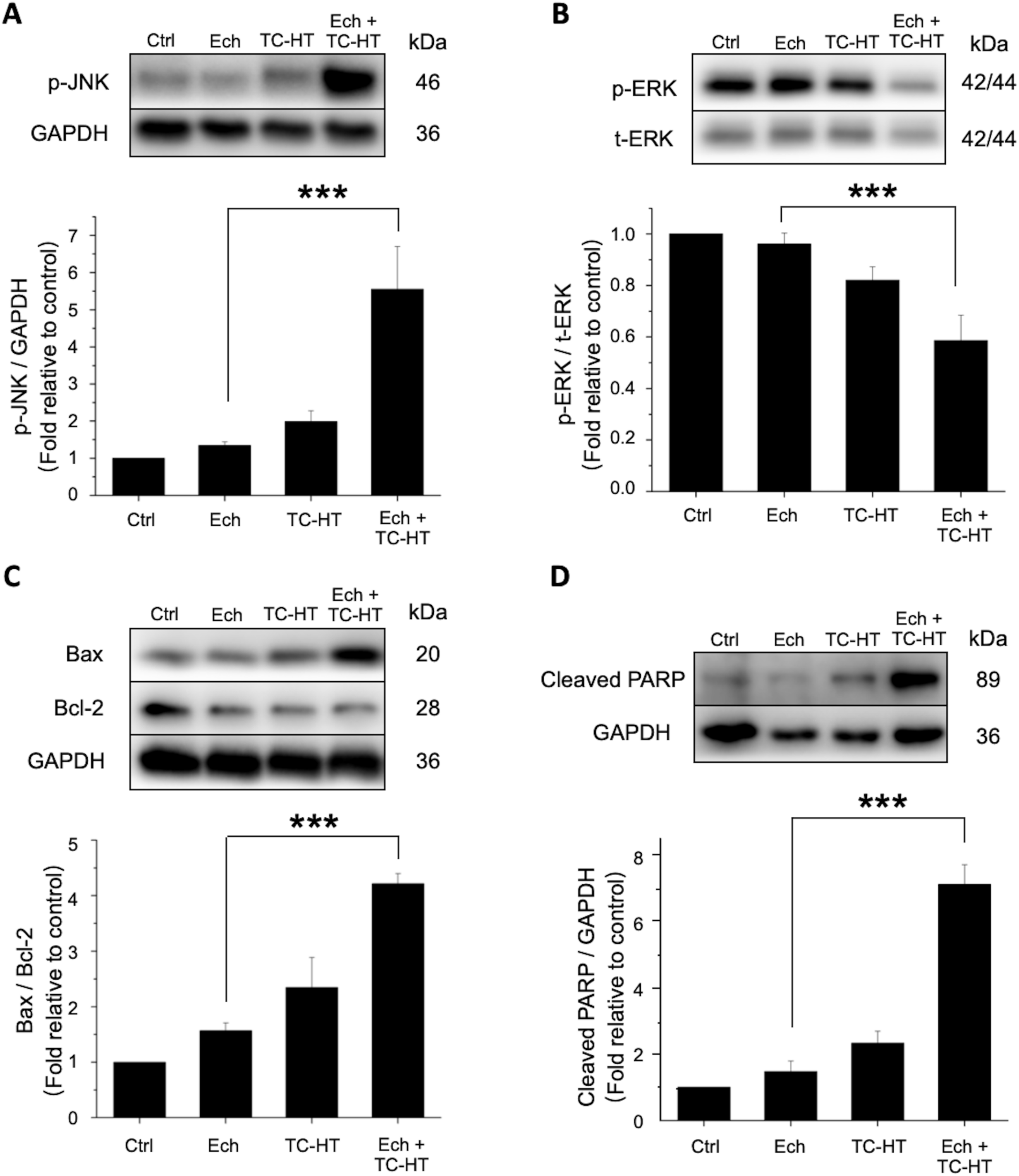
Effect of Ech combined with TC-HT treatment on survival- and apoptosis-related protein expressions in PANC-1 cells. The experiments were conducted in PANC-1 cancer cells treated with 20 µM Ech alone or in combination with TC-HT. Western blot analyses of p-JNK (A), p-ERK and t-ERK (B), Bax and Bcl-2 (C), and cleaved PARP (D) protein expressions. The expression levels of p-JNK and cleaved PARP were normalized to GAPDH, and the p-ERK was normalized to t-ERK. Bax/Bcl-2 ratio was calculated based on the Bax and Bcl-2 protein levels. Each relative expression level was compared with control and represented as fold relative to control. Data represent the mean ± standard deviation (n=3). Statistical significance was determined by one-way ANOVA followed by Tukey’s post-hoc test (***P < 0.001).

## Discussion

With the aim of investigating the anticancer effect of Ech and TC-HT on PANC-1 cells, the study finds that TC-HT strongly enhances the anticancer effect of Ech and causes death of PANC-1 pancreatic cancer cells significantly. Previous studies have proven the anticancer effect of Ech on some human cancer cell lines or animal models (23,24). Tang and co-workers, for instance, showed that Ech is capable of inhibiting breast cancer cells by suppressing the Wnt signaling pathway (23). Ye *et al*. demonstrated the anticancer effects of Ech in HepG2 cells and mouse model (24). With minimum harm to normal cells and tissues, natural products are still unsuitable for direct anticancer therapeutic application, due to insufficient efficacy and poor availability.

In this work, TC-HT was employed to significantly enhance the efficacy of natural product Ech, exhibiting a synergistic effect inhibiting cancer cell growth and inducing apoptotic cell death of PANC-1 pancreatic cancer cells. In the study, with only 20 µM Ech in combination with TC-HT, one can achieve the same cell inhibition effect as at 100 µM Ech alone. That is to say the Ech dosage required to achieve effective anticancer effect is greatly decreased to one fifth the original level, which also reduces damage to healthy cells. Such result is inspiring because one can achieve the same anticancer efficacy at much lower dosage. The enhanced specificity and efficacy of cancer therapeutic modalities and reduced side effect could further improve the survival rate of patients.

The study also finds that while TC-HT alone can suppress the expression level of MTH1, its combination with Ech further increases the inhibitive effect on MTH1 expression. Some studies have manifested that Ech inhibits tumor growth by induction of oxidative DNA damage or via specific signaling pathways (25–27). One crucial protein mediating the Ech-induced DNA damage is MTH1, which is responsible for sanitizing oxidized dNTP pools to prevent incorporation of damaged bases during DNA replication by, for example, converting 8-oxo-dGTP into 8-oxo-dGMP (28–30). Gad *et al*. showed that MTH1 is indispensable for cancer survival in several cancer cell lines (14). Moreover, previous study has reported that MTH1 depletion with interfering RNA or small molecule inhibitors may cause DNA damage and apoptotic cell death in various cancer cell lines (31). Since then, a growing number of MTH1 inhibitors have been developed in recent years (32–34). Furthermore, recent studies have provided more evidences that MTH1 inhibitors can delay tumor growth and inhibit metastasis both in vitro and in vivo (35,36), further upholding their potential in cancer treatment.

Ech is one of a few natural products found to have inhibitive effect on MTH1 (37). Dong *et al*. found that treatment of Ech inhibits MTH1 activity and increases DNA damage marker, inducing apoptotic cell death and cell cycle arrest in human cancer cell line, without harming normal cells (11). Our study demonstrates that TC-HT application significantly enhances Ech-induced MTH1 inhibition and DNA damage, as well as apoptotic cell death, while slashing the viability of PANC-1 pancreatic cancer cells to merely 20% of the control group. The outcome suggests that combined treatment has a similar in vitro efficacy as chemotherapy drugs. Moreover, with the assistance of TC-HT, it needs only 20 µM low dosage of Ech to attain the same cell inhibition efficacy as at 100 µM Ech alone.

Through western blot analysis, the study attributes the enhanced anticancer effect partially to the MTH1 protein inhibition triggered by the combination treatment. To the best of our knowledge, this is the first study demonstrating that Ech suppresses the MTH1 protein expression. The study also finds that TC-HT alone can effectively suppress MTH1 expression, its combination with Ech greatly enhances the effect. The DNA damage marker 8-oxo-dGTP also increases significantly in the combined treatment group, confirming the linkage between abrogated MTH1 and DNA damage. Dong *et al.* showed that Ech treatment could increase the cellular level of 8-oxo-dGTP, but the ROS level remained unchanged after 24 h treatment of Ech (11), perhaps due to the fact that Ech itself is also a potent antioxidant, thereby antagonizing the ROS level. Since cellular 8-oxo-dGTP is also generated by ROS, increase in cellular ROS will cause more 8-oxo-dGTP accumulation, creating more apoptotic signals. In the study, with the assistance of TC-HT, we find that the combination treatment can significantly increase not only 8-oxo-dGTP level but also ROS level, compared with Ech or TC-HT treatment alone. Furthermore, many studies have demonstrated that ROS accumulation could induce apoptosis by regulating the MAPK family protein expressions (38,39).

In our Western blot analysis, the results show that the combination treatment of Ech and TC-HT strongly activates p-JNK, while significantly inhibiting p-ERK. The combined treatment produces severe oxidative stress, which activates the apoptosis signal through weakening cytoprotective p-ERK and activating stress-related p-JNK proteins. In addition, the Bcl-2 family proteins play an important role in regulating apoptosis and survival activities. Particularly, Bcl-2 protein, a key apoptosis-regulation protein, is responsible for the sequestration of pro-apoptotic proteins such as Bax. It binds to pro-apoptotic proteins to prevent the oligomerization and pore formation in the mitochondrial membrane (40). Moreover, it has been proven that Bcl-2 is a critical apoptosis inhibitor which accounts for cancer chemoresistance (41,42). Therefore, several Bcl-2 inhibitors have been developed to target the Bcl-2 protein and found to be synergistic with chemotherapeutic drugs (43,44). In this study, the result shows that the combination treatment significantly increases the Bax/Bcl-2 ratio, which primes cancer cells for apoptosis via mitochondria-dependent apoptosis pathway. As the anticancer mechanism triggered by the combination treatment is multipronged, the synergistic effect is similar to combining MTH1, ERK, and Bcl-2 inhibitors, facilitating the development of new anticancer drugs. Given the concern over their effect and massive dosage in need, few herbal medicines have been applied in anticancer treatment. In this work, our study proposes the application of TC-HT as a sensitizer, significantly boosting the anticancer effect of Ech and thus slashing its dosage in need, facilitating the application of Ech in cancer treatment.

In conclusion, combined TC-HT and Ech treatment targeting multiple pathways is a novel cancer therapy with significant anticancer potential. The approach facilitates apoptotic cell death in PANC-1 cancer cells via MTH1 inhibition, ROS and 8-oxo-dGTP accumulations, and MAPK and Bcl-2 family protein regulations. Compared with Ech administration alone, the co-treatment with TC-HT not only boosts a higher efficacy treatment but also provides a safer and more comfortable medical procedure, capable of contributing to the study of cancer treatment.

## Acknowledgments

The authors would like to acknowledge the Technology Commons in College of Life Science of National Taiwan University for use of flow cytometry system. We would also like to thank the Molecular Image Core Facility of National Taiwan Normal University under the auspices of the National Science and Technology Council for use of Laser Scanning Confocal Microscope System (ZEISS LSM880 with airyscan)/[BIO002514].

## Funding

This work was supported by grants from National Science and Technology Council (NSTC 112-2112-M-002-033 to CYC) and Ministry of Science and Technology (MOST 110-2112-M-002-004 and MOST 109-2112-M-002-004 to CYC) of the Republic of China. The funders had no role in study design, data collection and analysis, decision to publish, or preparation of the manuscript.

### Availability of data and materials

The data generated in the present study may be requested from the corresponding author.

### Authors’ contributions

CYC conceived and supervised the study. WTC, YMC and CYC designed the study. WTC, YMC, GBL, YYK, HHL and CYC performed the experiments, and were involved in data analyses and interpretation. WTC, YMC and CYC wrote the manuscript. WTC, YMC and HHL confirm the authenticity of all the raw data. All authors read and approved the final version of the manuscript.

### Ethics approval and consent to participate

Not applicable.

### Patient consent for publication

Not applicable.

